# How to define, use, and interpret Pagel’s λ (lambda) in ecology and evolution

**DOI:** 10.1101/2023.10.10.561651

**Authors:** William D. Pearse, T. Jonathan Davies, E. M. Wolkovich

## Abstract

1

Pagel’s λ (lambda) is a critical tool in ecology and evolution for describing trait evolution, imputing missing species’ data, and generalising ecological relationships beyond their study system. Yet the interpretation of λ depends on context, and there are many misconceptions about metrics that are similar but not identical to λ. As an index of phylogenetic signal applied to continuous traits, λ typically (but not always) ranges between 0 and 1, and is a rate-independent measure of the degree to which closely-related species resemble one-another relative to a Brownian motion expectation. But this measure is biased by non-random species sampling—a common characteristic of ecological data—which also makes phylogenetic imputation of missing traits challenging. The λ estimated in regression models has little to do with the phylogenetic signal of measured traits and is better considered as either a statistical correction or a measure of the impact of unmeasured (latent) traits in the model. In other contexts, such as hierarchical models including intra-specific variation, λ is frequently confused with distinct metrics such as *h*^2^. We show how confusion in defining and using λ can mislead our interpretation of ecological and evolutionary processes.

**Open research statement:** No data were used or collected as part of this work.

## 2 Introduction

Species traits are fundamental to our understanding of community assembly, the functioning of ecosystems, and how species might respond to environmental change (Bolnick et al., 2011; Foden et al., 2013; Violle et al., 2007). There is, therefore, large interest in modelling trait correlations, the interactions between traits, and in estimating trait lability. Because ecological interactions both shape and are shaped by the evolution of traits, integrating eco-evolutionary feedbacks operating over both short (Fussmann et al., 2007) and long (Cavender-Bares et al., 2009) timescales can help advance trait research. Reflecting this reality, these is now widespread application of phylogenetic comparative methods in ecology and global change research (Ackerly, 2009; Cavender-Bares et al., 2012; Edwards et al., 2007).

One of the most common uses of phylogeny in ecology is to detect and interpret patterns in species’ traits. Phylogenetic signal—the tendency for closely related species to share similar trait values—violates statistical assumptions of independence in many ecological analyses and so requires statistical correction (Freckleton et al., 2002). Correcting for this non-independence was one of the first uses of phylogenetic comparative methods (Felsenstein, 1985). Perhaps one of the most widely estimated phylogenetic signal metrics is Pagel’s λ (Pagel, 1999), a phylogenetic scaling parameter that describes how the shared evolutionary histories of species explain patterns of similarity observed in data. Often Pagel’s λ (lambda) is implicitly treated as the only relevant phylogenetic metric, but it is actually the third of the three phylogenetic transformations introduced by Pagel (1999). The other Pagel transformations, *δ* and *κ*, represent alternative modes of evolution that capture shifts in evolutionary rates, either across the depth of the tree (*δ*) or at speciation events (*κ*), but λ is the more frequently used (Münkemüller et al., 2012).

Here we review the definition of Pagel’s λ which, by virtue of being an algorithmic transformation, has different interpretations in different statistical contexts. We attempt to resolve some of the resulting confusion by considering its usage as an estimate of the phylogenetic signal in a single trait, including the impact of missing data and incomplete sampling in its estimation, as a scaling parameter estimated in classical trait regression, and in the emerging field of flexible hierarchical models of species’ ecological responses to environmental change.

We highlight five key points: (1) when estimated on a single trait, λ measures whether trait evolution is consistent with Brownian motion, which is not the same as evolutionary rate or phylogenetic correlation. (2) Ecological assembly is non-random, and so estimates of signal in ecological communities should be treated with caution. (3) Imputing traits on the basis of phylogeny is fraught with uncertainty, even if taxonomic sampling is fair and random (which, unfortunately, it rarely is). (4) λ in Phylogenetic Generalised Least Squares models has essentially very little to do with the phylogenetic signal of the modelled traits, but is an important statistical correction and can provide insight into the broader drivers of trait relationships. (5)

New hierarchical methods offer exciting opportunities to estimate the evolution of species’ ecological responses, but Pagel’s λ cannot be estimated from variance ratios and thus is not the same as heritability (*h*^2^).

## 3 Defining Pagel’s λ

It is remarkably difficult to pinpoint exactly when Pagel’s λ was first defined, but the most common citation is to Pagel (1999) where λ is discussed in detail (the original citation and derivation of λ remain somewhat obscure). λ is broadly described as a multiplier of a phylogeny’s internal branch lengths, but is more precisely—and correctly—described as *a multiplier of the off-diagonal elements of the phylogenetic variance-covariance matrix*. For example, in a three-species phylogeny (as depicted in figure 1), the λ transformation is applied to the phylogenetic variance-covariance matrix (*V*_*phy*_) to produce a resulting variance-covariance matrix (Σ) as follows:

**Figure 1:**
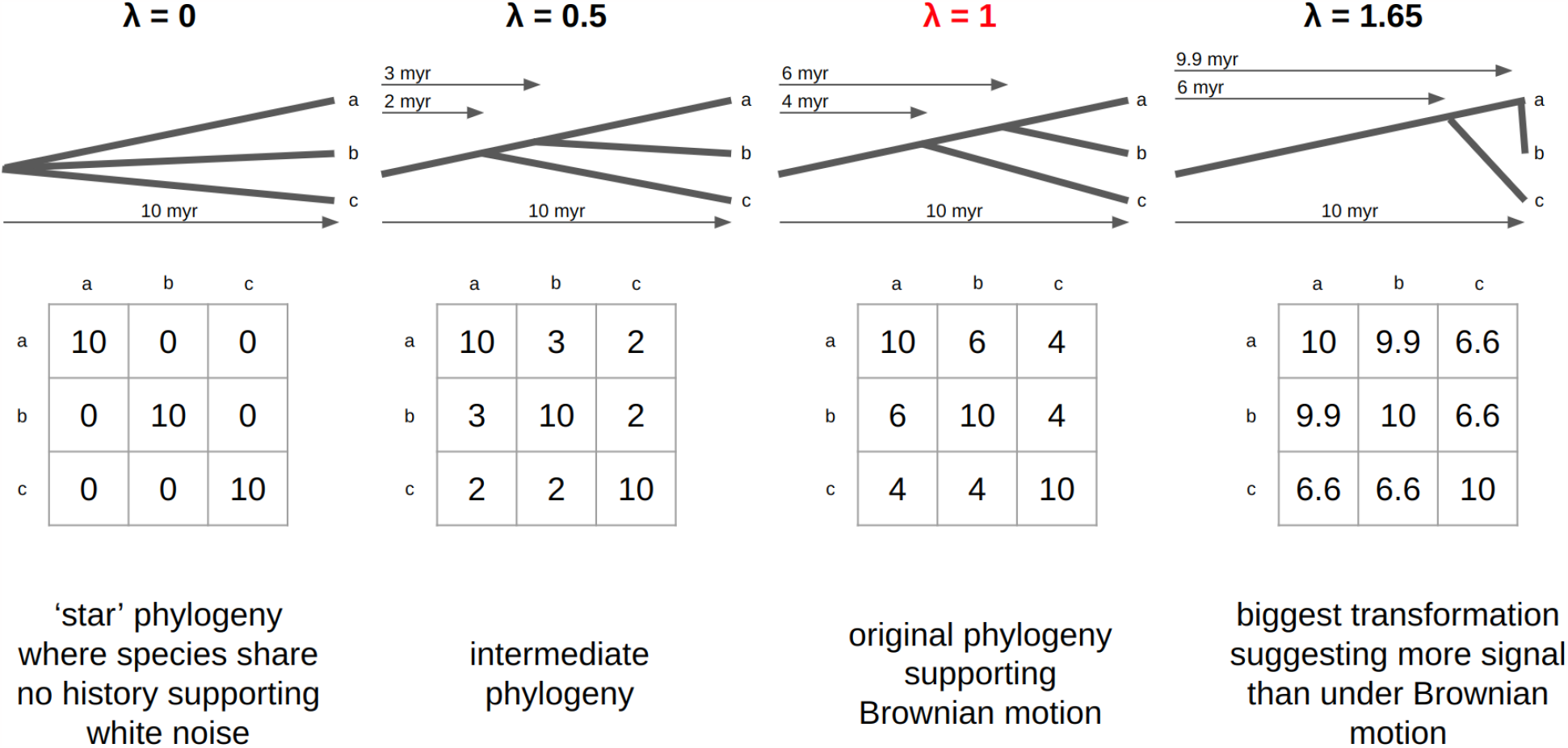
**Conceptual diagram of** λ. For each λ transformation (given at the top) the resulting phylogeny is shown below (with arrows to indicate branch lengths in millions of years), and the phylogenetic variance-covariance (*V*_*phy*_) matrix below that, and finally some comments on the transformation value below. Throughout, species’ names are indicated with letters (‘a’, ‘b’, and ‘c’). A λ of 1 leaves the tree unchanged (marked in red), while a λ of 0 transforms the phylogeny into a so-called ‘star phylogeny’ where essentially all phylogenetic information has been lost. The largest value of λ has been chosen to demonstrate the upper limit of λ in this phylogeny, where a further increase in λ would return off-diagonal elements (λ *· Σ*1, 2 and λ *· Σ*_2,1_) greater than one of the diagonal elements 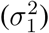, rendering *V*_*phy*_ improper. Conceptually, such a transformation reflects the impossibility of stretching the youngest node out beyond the present.

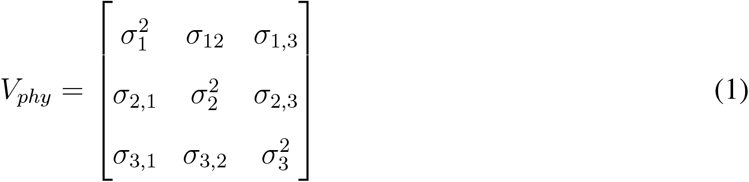

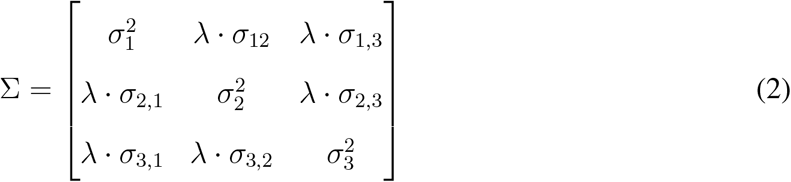

Where *σ* refers to (co)variances indexed by their subscripts, such that, for example, 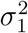 reflects the ‘diagonal’ variance unique to the first species, and *σ*_1,2_ the covariance shared by species 1 and 2. The variances (the diagonals of Σ) are equal to the root-to-tip distances for each species on the phylogeny and are unaltered by λ; the covariances reflect the shared branch-length between pairs of species.

λ was initially introduced to capture the degree of similarity among species as an index of phylogenetic signal in a measured trait: “*whether the phylogeny correctly predicts the patterns of covariance among species on a given trait*” (Pagel, 2002b). However, Pagel (2002a) also describes λ as a measure of the “*extent to which it is necessary when investigating trait evolution to take the phylogeny into account*”. With the rapid expansion of phylogenetic comparative methods following Felsenstein’s seminal paper on independent contrasts (1985), the latter usage has become widely adopted. Pagel’s λ was subsequently incorporated into the phylogenetic regression, popularised in Phylogenetic Generalised Least Squares (PGLS; Freckleton et al., 2002) as implemented in the *R* (R Core Team, 2021) package ‘*caper*’ (Orme et al., 2013).

Not all implementations or definitions of λ are the same: the *R* package *brms* (Bürkner, 2021), for example, defines it in terms of a variance ratio (equation 5; discussed in detail below). These varying definitions in part reflect the pace of development in the field of comparative methods, but also make it more difficult to compare estimates of λ. As we outline below, while it has been suggested that other formulations are equivalent, those equivalencies are, at best, controversial and, at worst, wrong; so we suggest avoiding such simplifications.

## 4 Describing the evolution of a single trait

Pagel’s λ is frequently used to describe the extent to which species’ trait values are correlated with phylogeny, whether past evolution has been slowed by some constraints, or whether a particular ecological force has impacted the evolutionary trajectory of the trait. In many cases these interpretations are misinformed.

Pagel’s λ measures phylogenetic signal, and uses Brownian motion as its marker of that signal. While there are alternative approaches for quantifying and testing for phylogenetic signal, or the related concept of phylogenetic niche conservatism, we do not explore them here as they have already been extensively reviewed and debated elsewhere (*e*.*g*., Blomberg et al., 2003; Diniz-Filho et al., 2012; Losos, 2008; Münkemüller et al., 2015; Münkemüller et al., 2012; Wiens et al., 2010).

When estimated for a single continuous trait on an ultrametric phylogeny with branch lengths proportional to time and tips (species) terminating at the present, λ represents the degree to which the phylogenetic distance between species corresponds to differences in their trait values (figure 1). Values of λ = 0 match to a white noise model of evolution and indicate no phylogenetic structure in traits: species’ phylogenetic relationships do not predict their trait similarities. This can also be represented by a ‘star’ phylogeny. Values of λ = 1.0 correspond to traits matching the assumptions of Brownian motion, in which trait evolution approximates a random (“drunkard’s”) walk through trait space. Typically λ = 1.0 is interpreted as evidence of strong phylogenetic conservatism in trait evolution, although the definition, and best measure of phylogenetic conservatism has been a matter of debate (Losos, 2008; Münkemüller et al., 2015). Phylogenetic signal and niche conservatism, however, are not the same (Losos, 2008; Wiens et al., 2010). When 0 *<* λ *<* 1, which is what we most commonly observe (we are unaware of a systematic review of such values, but see Harmon et al., 2010; Pennell et al., 2015), species’ traits appear intermediate between white noise and Brownian motion models. In such intermediate cases, traits are less similar amongst species than expected from their phylogenetic relationships under assumptions of Brownian motion, but closely related species are more similar to each other than expected by chance. λ is often sensitive to the presence of some degree of phylogenetic signal, but its confidence intervals are sufficiently wide that λ estimates are rarely precise (Münkemüller et al., 2012). A λ transformation is sometimes used in models of discrete (non-continuous) trait evolution [*e*.*g*., see *geiger::fitDiscrete* (Pennell et al., 2015)], where similar caveats apply to their use, and alternative statistics have been proposed to better capture evolutionary signal in binary traits (*e*.*g*., Fritz & Purvis, 2010).

Values of λ *>* 1.0 have occasionally been reported, but are controversial (Revell et al., 2008). While Pagel allowed λ to take such values, he explicitly stated that their interpretation was not defined (Pagel, 2002b), and many modern fitting routines do not permit them (Orme et al., 2013; Pennell et al., 2014). Biologically, a λ greater than 1 represents a stretching of internal nodes towards the present day (*i*.*e*., the pendant edges of the phylogeny). Such a transformation increases the evolutionary history that species share and so can be considered to be allowing for more phylogenetic conservatism than a Brownian motion model across a phylogeny would normally imply. Nonetheless, there is an upper bound to λ defined by the ratio of the total tree age to the age of the youngest node (figure 1). Framed statistically, as λ is a transformation of the phylogenetic covariance matrix, and—as no valid covariance matrix can have an off-diagonal element larger than its diagonals—the upper bound of λ is determined by the ratios of the largest off-diagonal and diagonal entries (see also Revell & Harmon, 2022). Empirically, at this λ upper-limit the youngest node is brought to the present day and so no higher λ values can be applied to the phylogeny because we measure traits in the present. Thus values *>* 1 are rarely fit in practical terms because that upper limit is idiosyncratic to each phylogeny, complicating comparison and interpretation among datasets.

Values of λ *<* 0 are not allowed, and would imply negative covariances such that trait values are predicted to be more similar between less closely related species. Negative auto-correlation is a regular feature of social networks (*e*.*g*. De Nooy, 2013), finance systems, (*e*.*g*. Koutmos, 1997) and spatial data (*e*.*g*. Griffith & Arbia, 2010), but has been less explored as a model of trait evolution. However, competitive character displacement and evolutionary convergence could both, in theory, give rise to negative covariances (*i*.*e*., traits that are over-dispersed relative to Brownian expectations; Davies, Cooper, et al., 2012). Such negative correlation has been termed ‘anti-signal’, but we have not been able to find a definitive first reference for it in that context. Nonetheless, negative patterns have been extensively studied in community phylogenetics (termed ‘phylogenetic over-dispersion’) as a pattern (Webb et al., 2002) and in phylogenetic generalised linear mixed modelling (Gallinat & Pearse, 2021; Ives & Helmus, 2011).

λ is a metric of the degree of fit with Brownian motion evolution, and so is independent of the rate of evolution (Revell et al., 2008) which is included as a separate parameter (*σ*^2^) in Brownian motion models. There are various metrics of the rate of trait evolution (*e*.*g*., the Felsen; Ackerly, 2009), but λ is independent from them. We provide an illustration of this in the supplementary materials (see also (Revell et al., 2008)). That a trait is consistent with Brownian motion evolution does not mean either that the trait has evolved under strong selection or that the trait has evolved in the absence of selection (Revell et al., 2008). Brownian motion could be consistent with both the evolution of a fitness-neutral trait (Losos, 2008) or strong fluctuating selection (Revell et al., 2008), and there is little way to separate the two. Notably, standard models of Brownian motion do not contain a parameter to indicate the strength of selection, and the most commonly-used macro-evolutionary model that does (Ornstein-Uhlenbeck or OU models; Butler & King, 2004) is, itself, only contentiously mapped onto macro-evolutionary process (Cooper et al., 2016; Pearse et al., 2018).

## 5 The dangers of missing species and biased sampling

In estimating λ we assume the underlying phylogeny is known without error, which is almost certainly not the case. Fortunately, Pagel’s λ is relatively robust to incomplete phylogenetic resolution and branch length errors (Molina-Venegas & Rodríguez, 2017), unlike some alternative metrics of phylogenetic signal such as Blomberg’s K (Davies, Kraft, et al., 2012). Nonetheless, λ estimated on an incorrect tree topology will necessarily also be incorrect. Here, we highlight a more pernicious but under appreciated source of error: the choice of species for an analysis. In the supplementary materials, we give code examples of the impacts of these well-studied issues.

Pagel’s λ was defined and derived under the expectation of a completely sampled, monophyletic dataset, and while λ tests are relatively robust to incomplete samples those samples should be representative of the true underlying phylogenetic distribution of trait values (Münkemüller et al., 2012). Because, Brownian motion is a noisy process it is possible, and indeed probable, that λ will differ when estimated on different species subsets (*e*.*g*. Rafferty & Nabity, 2017) even when the underlying evolutionary model is invariant. When species sets share ancestral lineages, each branch on the phylogeny can have only one true evolutionary history for a given trait; thus it would be a mistake to interpret such differences as reflecting divergent evolutionary histories. Estimates of λ derived from community data are even more challenging. Observed ecological assemblages rarely, if ever, contain the entirety of a monophyletic lineage: indeed, most ecological processes result in a non-random subset of species (often biased by ecological filtering on the traits of interest themselves). Differences in λ estimated from phylogenetically overlapping species sets might provide information on the process of community assembly (Prinzing et al., 2021) or reveal environment-by-trait interactions for spatially structured data (*e*.*g*. Davies et al., 2013). But non-random, ecologically filtered assemblages would not be expected to give reliable insight into the broader evolutionary forces describing trait evolution across the entire clade.

## 6 Phylogenetic imputation of missing trait values

When measured traits on a species set exhibit strong phylogenetic signal, it can be tempting to use phylogeny to impute trait values for unsampled species (Bruggeman et al., 2009). Phylogenetic imputation provides an approach for gap-filling traits datasets without the need for expensive and time consuming field work and trait measurement (Swenson, 2014). However, high λ in measured traits is insufficient to ensure high accuracy in imputed data, and the noisiness of the Brownian motion process makes predictions for species with few close-relatives particularly unreliable even for traits showing strong evolutionary conservatism (Molina-Venegas et al., 2018). In some cases, imputed values have been shown to be indistinct from noise (Molina-Venegas et al., 2023). Nonetheless, particularly when combined with correlated trait data, phylogenetically informed models can provide powerful approaches for interpolating missing values (Debastiani et al., 2021; Penone et al., 2014) and should be preferred over non-phylogenetic models. The simplest solution to handling uncertainty in imputed values is to ensure that errors associated with trait imputations are propagated forward into subsequent analyses. This includes both the uncertainty from the Brownian motion parameter (*σ*^2^) and its impact on the standard error of the estimate, as well as the fit of the Brownian motion model itself (*i*.*e*., model uncertainty).

## 7 Regressing two, or more, traits against one-another

Ecology has long been interested in trait correlations, using regression models to understand how they relate to each other. In the 1990s and early 2000s there was great debate over the importance of phylogeny in the regression of traits, especially as they relate to ecology (Harvey et al., 1995; Westoby et al., 1995), and that debate has somewhat reopened recently (Uyeda et al., 2018; Westoby et al., in press). While our purpose here is not to rehash that debate, a fair summary would be that if we are interested in generalising relationships we explore to other species and/or ecosystems, then an evolutionarily (phylogenetically) informed analysis is required to account for species’ non-independent evolution (Felsenstein, 1985; Harvey, Pagel, et al., 1991).

The development of phylogenetic comparative methods advanced rapidly over the 1990s and 2000s, with Phylogenetic Generalised Least Squares (PGLS; Freckleton et al., 2002), a generalisation of Felsenstein’s Phylogenetic Independent Contrasts (Felsenstein, 1985), now commonly employed to regress traits across species. These methods can reveal that observed trait correlations are the result of shared evolutionary history: this does not negate the obvious presence of a correlation, but rather suggests that the correlation may result from shared historical accident. To interpret λ in PGLS, we must first understand how the model is fit (which is detailed in Freckleton et al., 2002). There is no simple equation that will report the maximum likelihood λ value; instead, the model must be fit by maximising the likelihood (*L*) of the observed data given different model coefficients (including λ).

PGLS is a variant of Generalised Least Squares (GLS) and, as in all GLS models, the log-likelihood (*L*) function is:

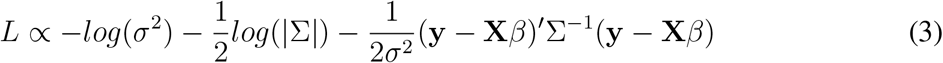

Where all terms are as already defined save the response variable (trait; **y**), an explanatory variable (trait(s); **X**), the model coefficients (*β*), and the variance (*σ*^2^; note that this is strongly linked to the Brownian motion definition in equation 2). PGLS is different from GLS when estimating λ only in that Σ is as defined in equation 2: its popularity and beauty stems from being able to make use of the existing body of statistical theory about GLS.

There is some confusion about what λ estimates in PGLS (see Uyeda et al., 2018, for an excellent discussion), perhaps in part because λ does not appear in the likelihood function (equation 3) but rather is implicit in the definition of Σ (itself defined in equation 2). Importantly, the λ estimated in a PGLS is *not* the phylogenetic signal of any of the traits involved in the regression. It is not only possible, but common, for two traits with phylogenetic signal to return an estimated λ of 0 when regressed against one another in PGLS (in the supplementary materials we give a demonstration of this surprising, but well-established, property). This is because a PGLS fits λ not just to the transformed matrix Σ but also to the residuals (species-specific errors; **y** *−* **X***β* in equation 3) in the model. However, PGLS can be used to estimate the phylogenetic signal of a trait if (and only if) the trait of interest is the response variable in the regression and the sole explanatory variable is the intercept. In this special case the residuals of the PGLS are equal to the variation in the response variable, and so the λ reported by a PGLS is the phylogenetic signal in that trait and can be interpreted as such.

One way to interpret PGLS is that it allows the phylogeny to capture some putative latent trait that affects the relationship between the directly modelled traits, and PGLS reveals the phylogenetic patterning to that latent trait. This latent trait may enhance or mask the estimated statistical relationship between the modelled variables that would be apparent in the absence of this confounding evolutionary pattern.

Phylogenetic regression (*sensu* Grafen, 1989) and PGLS are not the only ways to quantify multi-trait evolutionary information. Phylogenetically-informed versions of dimension-reduction techniques, such as ordination or principal component analysis (PCA), that leverage λ also exist.

In cases where the goal is to identify general patterns and groupings of traits (and there is no particular response variable), then phylogenetic redundancy analysis (RDA) and PCA (Revell & Harrison, 2008) are useful methods. While we do not consider those approaches here, we emphasise that λ estimated in a novel context (*e*.*g*., a PCA), will require careful reassessment, and may not always be interpreted analogously to λ in a PGLS, even if fit similarly as a transformation of the phylogenetic variance-covariance matrix.

## 8 Hierarchical models

With modern approaches (*e*.*g*. Hadfield et al., 2014; Ives & Helmus, 2011), and in particular the advent of flexible Bayesian methods (Uyeda & Harmon, 2014), we are no longer restricted to fitting models where phylogenetic information, such as λ, can only be accounted for in the residual error structure. These new approaches allow us to answer questions about how species’ responses to environment and other species have evolved, and are an exciting avenue for eco-evolutionary research (Gallinat & Pearse, 2021). There is a need for caution, however; as there are myriad links and generalisations that bind different hierarchical approaches it can be challenging to understand when they apply to λ.

The implementation of such hierarchical models is described elsewhere in greater detail (*e*.*g*. Gallinat & Pearse, 2021; Ives & Helmus, 2011), but in brief: hierarchical models can be fit where species’ coefficients (*e*.*g*., environmental responses or interaction coefficients) are drawn from distributions parameterised by phylogenetic covariance matrices that are, themselves, scaled by λ. We expand on the approach described by Ives (2019) to define such a covariance matrix (Σ), specifically:

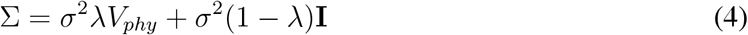

Where *σ*^2^ is the Brownian motion rate parameter and **I** the identity matrix (a matrix where the diagonals are all 1 and the off-diagonals are all 0). Equation 4 is essentially an algebraic re-arrangement of equations 1 and 2 to allow for a more efficient estimation of Σ that makes separating its phylogenetic (*V*_*phy*_) and non-phylogenetic (**I**) components more straightforward. In practice models may not be fit with precisely this formulation, more often employing some form of algorithmic shorthand or Cholesky decomposition to speed estimation, but the fundamentals are the same (Ho & Ane, 2014; Pearse et al., 2015).

It is possible and conceptually appealing to partition variance into phylogenetic and non-phylogenetic components, and thus derive the ratio of the phylogenetic component to the total variance, heritability (*h*^2^; see Hadfield & Nakagawa, 2010; Lynch, 1991):

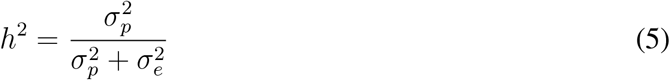

Where 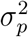 is a variance multiplied by a phylogenetic variance-covariance matrix (akin to *V*_*phy*_ in equation 4) during model fitting, and 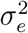 is the residual error variance across the entire model. While precise definitions for the terms within this equation depend on context, the spirit of it is always a variance ratio: variance associated with phylogeny 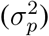 compared to the total variance in the model 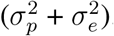.

Confusion can arise when comparing Pagel’s λ to such estimates of phylogenetic heritability from hierarchical models, and when trying to find equivalencies among them. There has been debate on the exact equivalency of λ and *h*^2^, since it is not possible to fit the two formulations above (equations 4 and 5) simultaneously, yet phylogenetic signal seems, intuitively, similar to the amount of variation in data explained by phylogeny. This intuition is almost always incorrect, however. Housworth et al. (2004) suggest the two metrics are mathematically equivalent; this cannot be the case since, as Freckleton et al. (2002) highlight, *h*^2^ is bounded between 0 and 1, but λ is not a ratio and can take values *>*1 as we discuss above. Confusingly, under strict assumptions of Brownian motion evolution, both λ and *h*^2^ are expected to be 1, and both are expected to be 0 for traits in the absence of phylogenetic structure (Freckleton et al., 2002). It thus seems reasonable to expect that for intermediary cases *h*^2^ and λ will also be similar, but this is not necessarily the case; those who, like us, have followed too closely advice from standard statistical software (*e*.*g*., Bürkner, 2021) need not panic, but simply recognise these different model formulations and carefully report the metrics calculated.

The variance partitioning approach has some limitations as it suggests separating coefficients that may be better represented as intertwined. For example, take a model of the form:

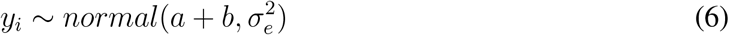

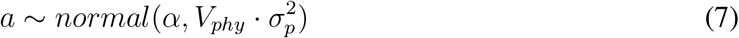

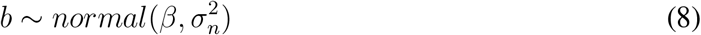

Where *y*_*i*_ are observations of some trait (*y*) across species (indexed by *i*) drawn from a normal distribution centred according to phylogenetic (*a*) and non-phylogenetic (*b*) components with some overall error variance 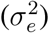. The species-level estimates are themselves drawn from a normal distribution centred at some mean (*α* or *β*) with a variance informed by a phylogenetic variance-covariance matrix (*V*_*phy*_) with some estimated phylogenetic variance 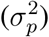 or independently 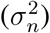. This kind of model (although most often used with replicate measurements of species and with additional explanatory variables) is advocated for in some ecological contexts (Bürkner, 2021), likely in part because PGLS is not well formulated to accommodate multiple observations across species (but see Freckleton & Rees, 2019; Joly et al., 2019). In such cases, *h*^2^ has been defined incorporating all three variance terms:

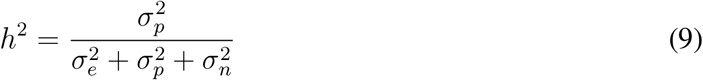

There are two problems with such a formulation. First, even if the original Lynch (1991) formulation maps onto Pagel’s λ equation 9, and the many diverse models it allows, it has not been formally confirmed to do so. Second, this formulation is statistically inefficient and potentially biased because it is co-estimating variance associated with species using two separate distributions (Bafumi & Gelman, 2007). The phylogenetic and non-phylogenetic terms are, in effect, competing for variance when they should be jointly fitting it.

The development of new statistical models that include phylogeny is exciting. Advances in model fitting and speed may allow for greater flexibility in estimating the evolution of species’ environmental responses and interactions (Gallinat & Pearse, 2021) and allowing for interactions with latent traits, represented by the phylogeny (Morales-Castilla et al., in review). However, along with such advances is a need to better understand the equivalencies of parameters estimated under alternative model formulations.

Lynch (1991) offered conceptual simplicity by estimating the fraction of variation that is explained by phylogeny. Pagel’s λ (1999) offers statistically efficiency, and maps nicely onto well-studied concepts such as Brownian motion evolution. Here we suggest an approach that combines the two: using a pooling metric, *ω* (named following Gelman & Pardoe, 2006), that captures the variance in model fit attributable to species-specific variation using Pagel’s λ transformation:

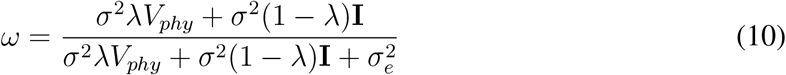

Where all other terms are as defined in equation 4 and above. This provides flexibility (additional hierarchical terms could be added and then included in the denominator) and speed of estimation (since all variance-covariance matrices would be estimated as outlined above in reference to equation 4). We suggest such a metric would be of particular use in forecasting responses in under-studied species, for which we may have phylogenetic information but limited ecological observations.

## 9 Summary

Pagel’s λ (lambda) is useful for measuring pattern in evolutionary process, and helps generalise findings beyond a particular ecological study system. We have discussed how λ is estimated, and the common pitfalls in its application and reporting. Specifically:

- λ is (usually) a number between 0 and 1 that measures phylogenetic signal, but care must be taken as it can exceed 1 and there are many different definitions of phylogenetic signal.
- When estimating signal in individual traits, λ is sensitive but imprecise, and non-random taxon sampling—common in ecology—can bias estimates.
- λ is independent of evolutionary rate, and the interpretation of differences in λ estimated on phylogenetically overlapping taxon sets is not straightforward.
- High λ is likely not a useful index of adaptive constraints to global change as it integrates across (macro)evolutionary processes operating over long timescales (often millions of years), and does not necessarily provide information on constraints to adaptation over short timescales.
- High λ is necessary but not sufficient for successful phylogenetic imputation of missing trait data.
- In classical regression modelling, λ allows us to account for latent traits that covary with phylogeny and potential statistical confounds, but it is not a panacea and is informative on the residual (latent) variation only.
- When estimated in a phylogenetic least squares regression, λ can be zero even when measured traits are phylogenetically structured, because λ is estimated using the residual errors.
- Hierarchical models are powerful tools to study the evolutionary history of ecological processes, but the foundational definitions and linkages among statistical approaches complicate interpretation of λ, and different derivations are not always interchangeable.
- Estimates of heritability (*h*^2^) derived from hierarchical models are often similar to Pagel’s λ, but are only equivalent to λ under a set of very limited situations.
- As we move forward with a new era of complex, flexible modelling, we must be careful to report precisely what we have calculated in our models, and not assume that equivalencies in one domain apply to another.

## Acknowledgements

David Orme and Nell Pates provided insightful comments on the manuscript.

