## Supplementary code for "How to define, use, and interpret Pagel’s λ (lambda) in ecology and evolution"

## Supplmement for: How to define, use, and interpret Pagel's \lam (lambda) in ecology and evolution
## Pearse, Davies, Wolkovich - 2023-09-22

## What follows are quick 'back of the envelope' demonstrations of
##   properties of Pagel's lambda that are already well-established in
##   the literature. For detailed and thorough demonstations of these
##   properties, please see the citations in the main text, but
##   otherwise what follows hopefully gives a flavour and is sufficient
##   to satisfy the curious non-specialist.

## IMPORTANT NOTE: This script sets random number seeds for reproducibility,
##                   so please restart R after using this script before conducting
##                   any kind of serious analysis.

##########################
## Packages ##############
##########################
library(caper)
library(geiger)
library(picante)
library(phytools)

## 1: Phylogenetic signal != evolutionary rate
## ("There are various metrics of the rate of trait evolution (e.g., the Felsen; Ackerly, 2009), but lambda is independent from them: the rate of Brownian motion is independent from an estimated lambda")
set.seed(123456)
tree <- sim.bdtree(seed=123456)
sigmas <- rep(10^seq(-2,1,length.out=30), each=3)
lambdas <- sapply(sigmas, function(x) phylosig(tree, sim.char(tree, x, root=0)[,,1], method="lambda")$lambda)
boxplot(lambdas ~ sigmas, main="Pagel's lambda plotted against rate of evolution (sigma)\n(n=3 for each sigma estmate)", ylim=c(0,1))
## ... as the rate of Brownian motion evolution changes across three orders of magnitude (sigma; horizontal axis), the estimated lambda is essentially unchanged.

## 2a: Taxon selection biases lambda
## ## ("Here, we highlight a more pernicious but under appreciated source of error: the choice of species for an analysis")
set.seed(123456)
tree <- sim.bdtree(seed=123456)
trait <- sim.char(tree, .1, root=0)[,,1]
phylosig(tree, trait, method="lambda") # In full dataset, lambda is accurately estimated to be 1
rnd.sampled <- trait[sample(1:100, 50)]
phylosig(tree, rnd.sampled, method="lambda") # With random sampling, lambda is still accurately estimated
bias.sampled <- trait[trait>mean(trait)]
phylosig(tree, bias.sampled, method="lambda") # With biased sampling, lambda is under-estimated
strong.bias.sampled <- trait[trait>(mean(trait)+.5)]
phylosig(tree, strong.bias.sampled, method="lambda") # With strong biased sampling, signal is not detected

## 2b: Taxon selection biases trait imputation
## ## (follows directly from the above)
rnd.estimated <- phyEstimate(tree, rnd.sampled)
bias.estimated <- phyEstimate(tree, bias.sampled)
par(mar=c(5.1, 5.1, 2.1, 4.1))
plot(rnd.estimated$estimate ~ trait[rownames(rnd.estimated)], axes=FALSE, xlab="True values", ylab="Randomly sampled imputation", pch=20)
points(bias.estimated$estimate ~ trait[rownames(bias.estimated)], pch=20, col="red")
abline(0, 1, lty=3, col="grey30")
axis(4, col="red", col.axis="red")
mtext(text="Bias sampled imputation", side=4, col="red", line=2)
axis(1);axis(2)
cor.test(rnd.estimated$estimate, trait[rownames(rnd.estimated)])
cor.test(bias.estimated$estimate, trait[rownames(bias.estimated)])
## ... These plots show how the sampling method for species (random in
##       black, biased in red, both plotted on vertical axis) affects
##       the accuracy of imputed traits (true values plotted on
##       horizontal). Note that the correlation coefficient between
##       the biased imputations is negative and absolutely small,
##       despite a relatively large sample size (50 species).

## 3 : PGLS lambda != signal in traits
## ## ("It is not just possible, but common, for two traits with phylogenetic signal to return an estimated \lam of 0 when regressed against one another in PGLS")
set.seed(123456)
tree <- sim.bdtree(seed=123456)
explanatory <- sim.char(tree, 1, root=0)[,,1]
response <- explanatory + rnorm(100)
c.data <- comparative.data(tree, data.frame(explanatory, response, species=names(explanatory)), species)
summary(pgls(response ~ explanatory, data=c.data, lambda="ML")) # PGLS lambda = 0
phylosig(tree, explanatory, method="lambda") # Explanatory trait lambda = 1
phylosig(tree, response, method="lambda") # Response trait lambda = 0.86
## ... => PGLS lambda can be 0 when traits show signal
